# Early and late RNA eQTL are driven by different genetic mechanisms

**DOI:** 10.1101/2025.02.24.639351

**Authors:** Saori Sakaue, Accelerating Medicines Partnership®: RA/SLE Network, Soumya Raychaudhuri

**Affiliations:** Center for Data Sciences, Brigham and Women’s Hospital, Harvard Medical School, Boston, MA, USA; Divisions of Genetics and Rheumatology, Department of Medicine, Brigham and Women’s Hospital, Harvard Medical School, Boston, MA, USA; Program in Medical and Population Genetics, Broad Institute of MIT and Harvard, Cambridge, MA, USA; Department of Biomedical Informatics, Harvard Medical School, Boston, MA, USA

## Abstract

Understanding the genetic regulation of RNA abundance is essential to defining disease mechanisms. However, conventional expression quantitative loci (eQTL) studies quantify RNA molecules across the transcript lifecycle. While most eQTL likely affect transcription by altering promoter or enhancer function within the nucleus, it is also possible that they modulate any processes after transcription, including chemical modifications and RNA stability in the cytosol. To elucidate distinct eQTL mechanisms of early versus late RNA, we compared eQTL from mature cellular RNA and nascent nuclear RNA in the brain and the kidney. Across tissues, we identified different causal variants for cellular and nuclear eQTL for the same *eGene*. Cellular eQTL were enriched in transcribed regions (*P*=3.3×10^-126^), suggesting the importance of post-transcriptional regulation. Conversely, nuclear eQTL were enriched in distal regulatory elements (*P*=7.0×10^-32^), highlighting the role of DNA transcriptional regulation. For example, we identified stop-gain eQTL variants likely acting through nonsense-mediated decay in cellular eQTL that had no effect in nuclear eQTL. Cellular eQTL were enriched for loci with multiple causal variants in linkage disequilibrium within the transcribed regions, where they may in concert affect RNA stability. We also identified examples of nuclear eQTL variants within enhancers that had no effect in cellular eQTL. We show that such eQTL (e.g., *TUBGCP4*) sometimes uniquely colocalize with disease alleles (schizophrenia). This study reveals key differences in the genetic mechanisms of cellular and nuclear eQTL.

## Introduction

Across their entire lifecycle, RNA molecules are under tight temporal and spatial regulation. In eukaryotes, promoters and enhancers in genomic DNA regulate RNA transcription in the nucleus^1^. The nascent transcript is then capped, spliced and polyadenylated to form mature RNA. Mature RNA molecules are exported to the cytoplasm, where RNA-binding proteins (RBPs) make chemical modifications and modulate structural stability, subcellular localization, translation efficiency, and degradation^2,3^. Safeguarding mechanisms for each step, such as nuclear retention in the nucleus and nonsense mediated decay (NMD) in the cytosol, prevent aberrant RNA from being translated^4^. The breakdown of high-fidelity RNA processing may contribute to disease^5^.

The effect of complex disease alleles on RNA abundance has been extensively cataloged through expression quantitative loci (eQTL) studies^6,7^. While eQTL studies have emerged as a valuable resource to interpret human disease alleles, most disease alleles from genome-wide association studies (GWAS) do not colocalize with eQTL loci^8–10^. Notably, most previous eQTL studies used RNA-seq to quantify transcripts from the entire cell. A majority of causal eQTL variants fine-map to gene promoters, while a minority localize to enhancers^6,11,12^. In either case, the emerging consensus is that eQTL variants regulate the transcription from the DNA by altering the sequence of DNA regulatory elements.^6^ However, genetic variants may affect any process after transcription, including post-transcriptional modifications essential for mature RNA stability. Indeed, molecular QTL studies have identified many loci that affect post-transcriptional processes, such as splicing^13,14^, alternative polyadenylation^15,16^ and N^6^-methyladenosine (m6A)^17,18^.

Here we assert that single-nucleus (sn)RNA-seq predominantly measures nascent RNA molecules, whereas single-cell (sc)RNA-seq or bulk RNA-seq measures the entirety of RNA molecules in the cells. The majority (70-90%) of cellular RNA captured by bulk cellular RNA-seq comprised cytosolic RNAs^19^. These different RNA profiling technologies can be used to assess the presence of distinct genetic determinants of expression at different timepoints in RNA lifecycle. Therefore, we conducted comparative analyses of eQTL variants from nuclear and cellular RNA. We obtained nuclear and cellular RNA-seq data in matched human tissues and conducted *cis*-eQTL analyses separately for each of the two modalities. We fine-mapped putative causal eQTL variants and identified systematic differences in the genomic locations of nuclear and cellular eQTL. We uncover how genetic regulation varies across RNA lifecycles and subcellular localizations.

## Results

### eQTL analyses with distinct RNA lifecycles

In this study (**Figure 1a**), we analyzed two tissues: (i) brain samples from dorsolateral prefrontal cortex (DLPFC) from participants in the Religious Order Study (ROS) and the Memory Aging Project (MAP)^20,21^ for discovery and (ii) kidney samples from lupus nephritis and control participants in the Accelerating Medicines Partnership (AMP) RA/SLE^22,23^ for replication **(Figure 1b)**. The DLPFC study was larger, consisting of cellular bulk RNA-seq (*n*_samples_=1,092) and snRNA-seq (*n*_samples_=424, *n_nuclei_*=1,562,635), while the kidney dataset offered direct comparison between scRNA-seq (*n*_samples_=179, *n_cells_*=547,796) and snRNA-seq (*n*_samples_=50, *n_nuclei_*=144,385). We first mapped *cis*-eQTL separately in both modalities and statistically fine-mapped causal variants for *eGene*s using approximate Bayesian factor (ABF) analysis^24,25^ (**Methods**). For each shared *eGene* in both nuclear and cellular RNA eQTL (n-eQTL and c-eQTL), we assessed whether causal variants of n-eQTL and c-eQTL colocalize^26,27^, and whether causal variant locations were systematically enriched within specific functional genomic annotations in one modality relative to the other (**Methods**).

**Figure 1.**
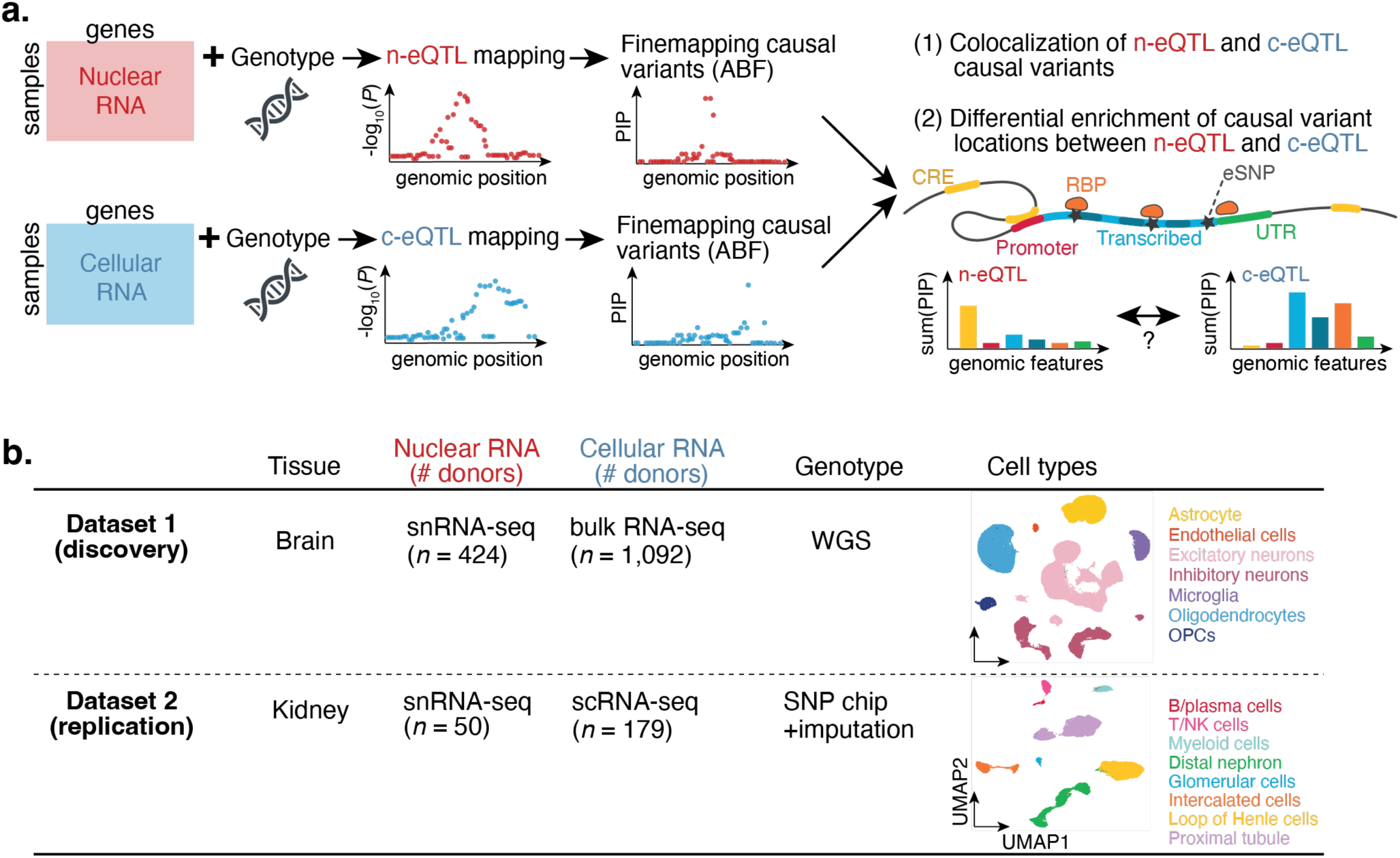
Overview of the analyses and datasets. **a**. Analytical framework. We mapped *cis*-eQTL of nuclear RNA (red) and cellular RNA (blue) in matched tissues, fine-mapped causal variants, and investigated systematic differences in causal variant location between nuclear eQTL (n-eQTL) and cellular eQTL (c-eQTL). **b**. Discovery dataset in the brain and replication dataset in the kidney. The cell type column depicts UMAP representation of the snRNA-seq and/or scRNA-seq dataset, colored by annotated cell types in the original study. WGS; whole genome sequencing. SNP; single nucleotide polymorphism. OPCs; oligodendrocyte progenitor cells.

In the discovery brain dataset, we identified 9,618 and 9,056 *eGene*s for nuclear snRNA-seq and cellular bulk RNA-seq, of which 5,632 were shared. We confirmed high consistency of our n-eQTL effect estimates with the original publication (Pearson’s *r*=0.98 in excitatory neurons; **Supplementary Figure 1**). For the 5,632 shared eGenes, the effect sizes for the n-eQTL lead SNP partly correlated with those for the c-eQTL (Pearson’s *r*=0.72; **Figure 2a**). We then tested if the n-eQTL and c-eQTL shared the same causal variant^26,27^ (**Methods**). Surprisingly, 1,835 of the shared *eGene*s (33%) had distinct causal variants (posterior probability (PP) for hypothesis 3 [H3]>0.7 in coloc). In contrast, 3,071 *eGene*s (55%) confidently had the same causal variants (PP(H4)>0.7; **Figure 2b**). In a comparison of two similarly sized bulk RNA-seq datasets (c-eQTLs) of the prefrontal cortex, PsychENCODE^28^ (*n*_samples_=1,387) and ROSMAP, we observed higher correlation of the lead SNP effect sizes (Pearson’s *r*=0.81) and colocalizing eGenes (73% with PP(H4)>0.7; **Supplementary Figure 2a** and **2b**). This suggests that the distinct causal variants for n-eQTL and c-eQTL are more likely driven by biological differences in the regulation of nuclear and cytosolic RNA, rather than by platform-specific or statistical fluctuations.

**Figure 2.**
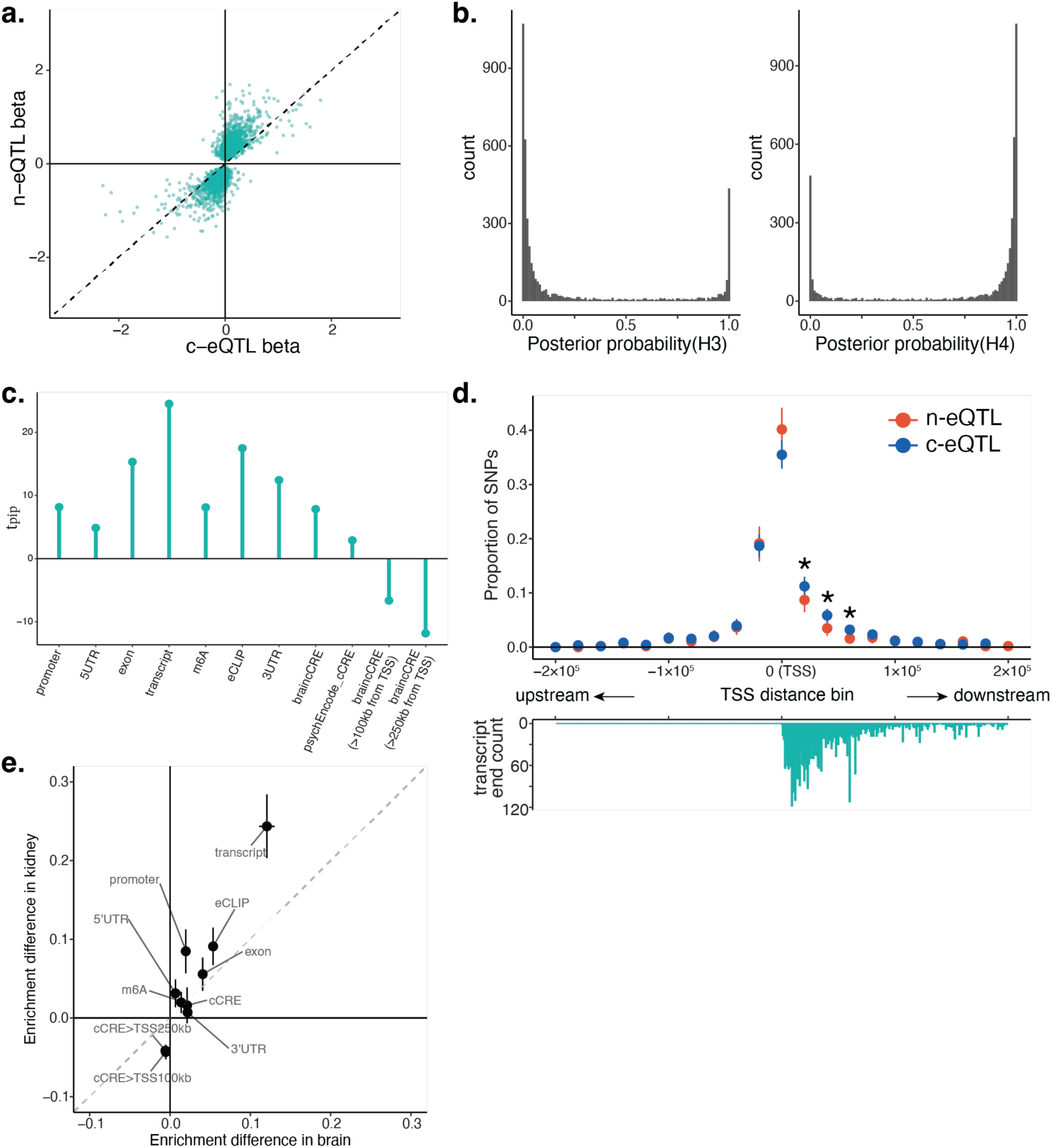
Distinct location of causal variants between nuclear eQTL and cellular eQTL. **a**. Scatter plot for beta coefficients for the same eSNP-eGene pairs between c-eQTL (x-axis) and n-eQTL (y-axis) in the brain dataset. Each dot represents one eSNP-eGene pair, and eSNPs were selected based on n-eQTL statistics. **b**. Histogram of coloc posterior colocalization probability for H3 (left; distinct causal variant between c-eQTL and n-eQTL) and H4 (right; shared causal variant between c-eQTL and n-eQTL). **c**. *t*_*pip*_ statistics comparing enrichment of c-eQTL and n-eQTL causal variant locations in the brain dataset. **d**. Proportion of SNPs within each genomic bin among all eSNPs with PIP > 0.7 for n-eQTL (red) and c-eQTL (blue). Stars above the bins indicate significant deviation from equal proportion between n-eQTL and c-eQTL (two-tailed chi-squared test *P*<0.05). **e**. Comparison of enrichment statistics differences in *t*_*pip*_ analyses between brain dataset (x-axis) and kidney dataset (y-axis). Bars indicate 95% confidence intervals.

### Distinct genomic locations of causal variants for nuclear and cellular eQTL

After fine-mapping, we examined the genomic features harboring eQTL causal variants. We derived the posterior inclusion probability (PIP) of variants in *cis* regions of a shared *eGene* and calculated enrichment of variants within the following genomic annotations (**Methods**): promoter, candidate *cis*-regulatory elements (cCREs), exonic region, transcribed region, 3’UTR, 5’ UTR, RBP binding sites by eCLIP assay^29^, and m6A modification sites^17^. Consistent with prior observations, we observed known enrichment in promoters, exonic regions and 5’ UTRs^30^ for both n-eQTL and c-eQTL (**Supplementary Figure 3**).

We then assessed if there was differential enrichment between nuclear and cytosolic RNA causal variants. We defined a differential enrichment statistic *t*_*pip*_ for each of the genomic annotations to test preferential enrichment for causal variants in n-eQTL or c-eQTL (**Methods**). Briefly, for each modality we took the aggregate of PIPs for variants within each annotation for an *eGene*; then we used a paired T-test to compare these values between n-eQTL and c-eQTL across *eGene*s. A positive *t*_*pip*_ indicates that the annotation has stronger enrichment for c-eQTL causal variants, whereas a negative *t*_*pip*_ indicates enrichment for n-eQTL.

Applying *t*_*pip*_ to 5,632 eGenes in the brain data, we found that c-eQTL causal variants are most strongly enriched for the transcribed regions (*t*_*pip*_=24.5, *P*=3.3×10^-126^), followed by exonic and RBP binding sites by eCLIP (**Figure 2c**). Given that chemical modification on the transcribed RNA and RBP binding sites affect folding, stability and decay of RNA in the cytosol^31^, this enrichment likely reflects genetic mechanisms that disrupt post-transcriptional modifications. Consistent with this hypothesis, the significant c-eQTL enrichment in exons and RBP binding sites remained significant even after conditioning on observed enrichment within transcribed regions (**Supplementary Figure 4**, **Methods**). Intronic regions, which in principle are spliced out and not exported into the cytosol, had instead enrichment for n-eQTL variants (*t*_*pip*_=-10.6, *P*=7.9×10^-26^ in conditional test; **Supplementary Figure 4**). Investigation of each of the 168 RBPs individually was underpowered, but we found that the majority of RBPs had positive *t*_*pip*_ indicating c-eQTL enrichment (**Supplementary Figure 5**). *RPS10* had significantly negative *t*_*pip*_ enriched for n-eQTL, and *DDX47* had nominally significant negative *t*_*pip*_ (**Supplementary Figure 5**). *RPS10* and *DDX47* had reported subcellular localization at granular component at nucleolus and nucleus, respectively, but not in the cytosol^32^. Among all tested annotations, we observed the most significant negative *t*_*pip*_ at candidate *cis*-regulatory elements (cCREs) more than 250 kb distant from transcription start site (TSS) (*t*_*pip*_=-11.8, *P*=7.0×10^-32^; **Figure 2c**), reflecting n-eQTL enrichment in distal enhancers. This might reflect weak enhancer’s regulatory effect at the transcriptional level within the nucleus that are more likely to be detected in the nucleus but buffered in the cytosol.

To support these observations, we examined the location of high-confidence eQTL variants (PIP>0.7) relative to the TSS. We observed significant differences in causal variant localization between n-eQTL and c-eQTL in the regions immediately downstream (3’) of the TSS corresponding to transcribed regions, with c-eQTL having more variants downstream of the TSS implicating transcribed regions (chi-squared *P*<0.05, **Figure 2d**; **Methods**). We confirmed this asymmetric distribution of causal variants (i.e., more causal variants downstream of TSS than upstream) in c-eQTL with a skewness statistic based on the third moment of the signed distances (**Methods**)^33,34^. A positive third moment indicates a right skewed distribution (downstream of TSS) and a negative third moment indicates the left skewed distribution (upstream of TSS). We observed the downstream skewness for c-eQTL (skewness=0.535 in c-eQTL and −0.406 in n-eQTL).

We considered that the observed differential enrichment might be driven by tissue or dataset specific effects. Hence, we replicated our findings in an independent kidney dataset with more matched profiling: snRNA-seq and scRNA-seq (**Figure 1b**). In this study, the snRNA-seq samples are a subset of the scRNA-seq samples. Despite different tissues and smaller sample size, we observed highly concordant *t*_*pip*_ with the discovery brain dataset across 248 kidney *eGene*s, with the most positive *t*_*pip*_ at the transcribed region (*t*_*pip*_=11.7, *P*=2.1×10^-25^), and the most negative *t*_*pip*_ at distal cCREs (*t*_*pip*_=-15.2, *P*=3.5×10^-37^; **Supplementary Figure 6a**). The mean differential effect estimates were highly correlated (Pearson’s *r*=0.95, *P*=2.6×10^-5^; **Figure 2e**).

Since the nuclear studies were smaller than the cytosolic studies, we considered the possibility that power differences in eQTL discovery and fine-mapping may influence the *t*_*pip*_. We performed secondary analyses (i) by statistically simulating equally powered n- and c-eQTL for the brain dataset in which raw expression data is unavailable for bulk RNA-seq and (ii) by using fully matched individuals between snRNA-seq and scRNA-seq for the kidney dataset. We replicated the same trend in *t*_*pip*_ across genomic annotations for both approaches (**Supplementary Figure 6b** and **6c**). We also confirmed that matching cell type proportions between snRNA-seq and scRNA-seq did not significantly change the observed enrichment (**Supplementary Figure 6d**). We concluded that the observed systematic differences in causal variant locations between n-eQTL and c-eQTL are pervasive across human tissues.

### Distinct biological mechanisms regulating RNA abundance in nuclear and cellular eQTL

We investigated the regulatory mechanisms explaining differences in causal variant locations between n- and c-eQTL by focusing on specific *eGene*s. First, we observed five eQTL variant (PIP>0.5) of stop-gain high-confidence loss of function (LoF) variants^35^ causing decreased gene expression in the brain. These variants still yield full transcript, but the resulting RNA with premature translation termination codons (PTCs) is degraded by NMD, preventing the production of harmful truncated proteins^36^. For example, in *VWA3B* locus, a highly-likely causal variant rs62154921 is a high-confidence LoF variant creating a stop codon (PIP=0.96). The stop-gain allele T significantly decreases the *VWA3B* expression in c-eQTL (beta=-0.91, nominal *P*=7.4×10^-35^), which is consistent with NMD. In contrast, the same variant rs62154921 showed low causal probability (PIP=3.0×10^-4^) in n-eQTL (**Figure 3a**). In the replication kidney dataset, there were no stop-gain variants in *eGene*s. There was one frameshift LoF insertion (rs66949844), again in c-eQTL with marginal probability of causality (PIP>0.1), which decreased the expression of *SIGLEC12*. Of these six LoF variant-eGene pairs, only one variant-gene pair (rs35233100 in *MADD*) was identified as significant both in c-eQTL and n-eQTL, with the remaining pairs identified solely in c-eQTL. NMD events occur in newly synthesized mRNAs as they exit the nuclear pore complex and simultaneously translated^37^. A previous study clarified that NMD occurred within 5–56 seconds of entering the cytoplasm^38^, which corresponds to the pioneer round of translation^39^. An exception has been reported in limited genes, in which NMD occurs after being fully released into the cytoplasm^40^, but none so far within the nucleoplasm. Therefore, our observation of NMD acting predominantly in c-eQTL is consistent with the previously demonstrated cytosolic localization of NMD, while one exception in n-eQTL might be explained by contamination of cytosolic RNA or ambient RNA^41^ into the extracted nuclei.

**Figure 3.**
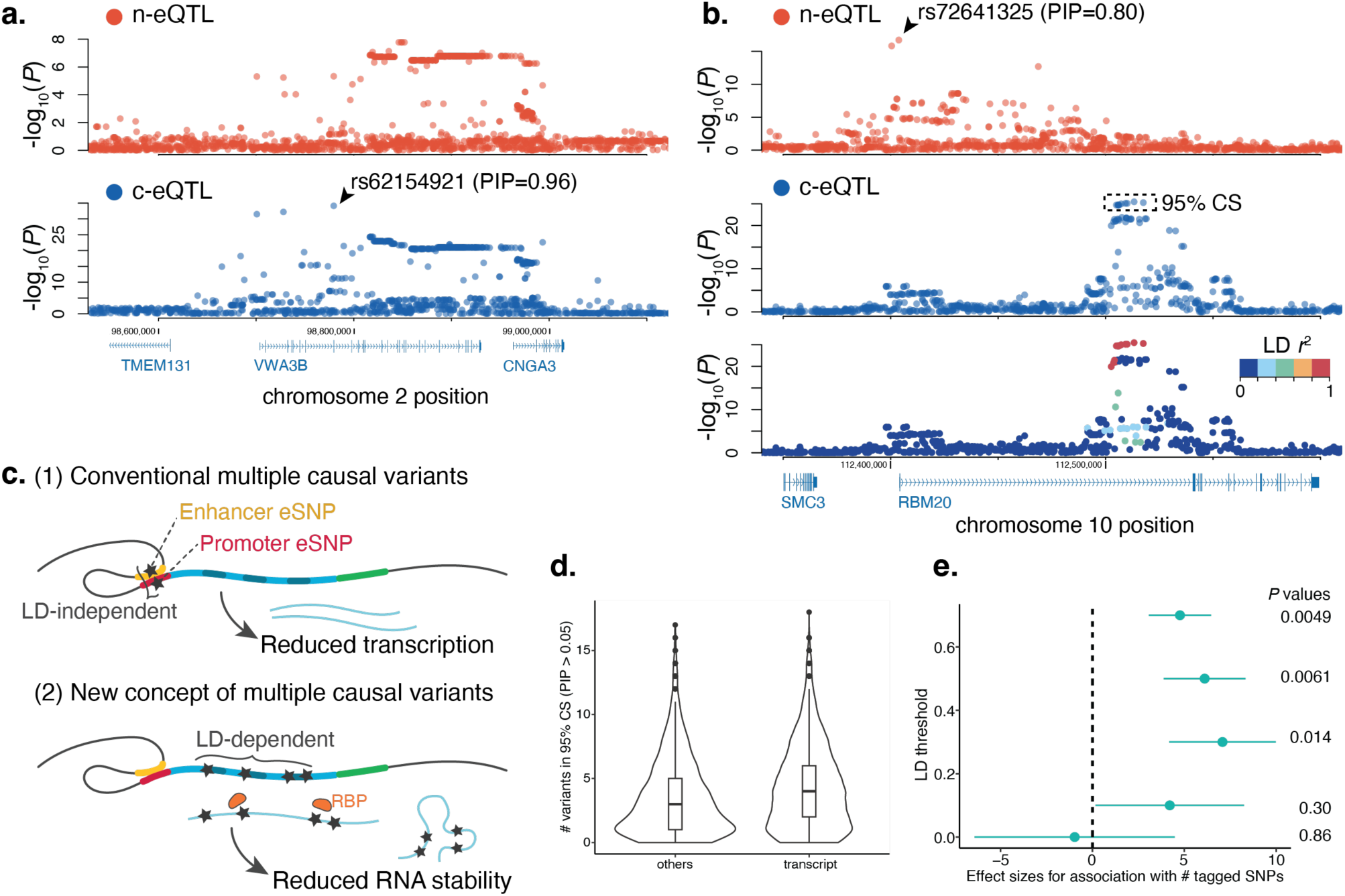
Different causal variants between nuclear and cellular eQTL reveal distinct biology. **a**. Stop-gain causal eQTL variant only observed in c-eQTL in the *VWA3B* locus. Top plot (red) and bottom plot (blue) depict regional association statistics for the n-eQTL and c-eQTL, respectively, at this locus. [Note: no variants with PIP > 0.05 in n-eQTL; 894 variants in 95% CS] **b**. distinct causal eQTL variant between n-eQTL and c-eQTL in the *RMB20* locus. Top plot (red) and middle plot (blue) depict regional association statistics for the n-eQTL and c-eQTL, respectively. The bottom plot depicts regional association statistics for the c-eQTL colored by LD *R*^2^ values as shown in the right upper color scale from the lead variant. **c**. Schematic representation of conventional and new concept of multiple causal variants for eQTL. **d**. Violin plots indicating the distribution of the number of variants with PIP > 0.05 within 95% credible sets for *eGene*s with causal variants within transcribed regions (right) and all the other eGenes (left). **e**. Beta coefficients (circles; x-axis) and standard error (bars) from the association between the LD score (i.e., sum of the LD *R*^2^ of *cis*-variants that had greater values of LD *R*^2^ than a sliding LD threshold (y-axis); **Methods**) and whether the *eSNP* is in the transcribed region or not.

Intriguing examples of distinct causal variants between n-eQTL and c-eQTL highlighted transcriptional vs. post-transcriptional regulation. There were 90 *eGene*s in the brain dataset with different causal variants between n-eQTL and c-eQTL with causal probability in transcribed regions <0.1 in n-eQTL but >0.7 in c-eQTL. Of them, 9 *eGene*s in n-eQTL had instead regulatory variants with causal probability in cCRE>0.7. *RBM20* is one such example (**Figure 3b**). In n-eQTL, we fine-mapped a single causal variant rs72641325 within cCRE proximal to the TSS (PIP=0.80). However, in c-eQTL, we identified a different 95% credible set of nine causal variants all located in the transcribed region. None of these variants individually had PIP>0.3 but together accounted for 95% of causal probability. Four variants in this credible set were located within the RBP binding sites (FUS, SAFB, RBFOX2, and KHDRBS1)^29^, suggesting functions in post-transcriptional modification and RNA structure. Moreover, these nine variants were in tight LD (*R*^2^>0.97; **Figure 3b**), indicating a shared haplotype with functional consequences on RNA structure. Such examples motivated us to consider the possibility that in certain instances multiple causal variants may drive eQTL and complex trait effects.

### New concept for multiple causal eQTL variants

Assuming a single causal variant or multiple causal variants in a locus has important implications for statistical fine-mapping approaches in eQTL and GWAS^42,43^. One approach to identify multiple causal variants is to use iterative conditional analysis, where a lead variant is identified and adjusted for to find new lead variants^44–49^. More recently developed methods, including SuSiE^50^, assume and create multiple ‘credible sets’ that are independent and each contains a subset of variants that explain each of the independent ‘single effects.’ Both approaches implicitly assume that we need to nominate and distinguish a true biologically causal variant from those that are non-functional and merely tagged by LD, regardless of the number of causal variants or credible sets in the locus. Functional genomic annotation has been used to assist this effort^30,51–54^. Now, our observation of *eSNP* haplotypes in the transcribed regions might propose a specific scenario that challenges this implicit assumption: correlated multiple variants tagged by LD (**Figure 3c**) may in concert influence RNA stability. In this scenario, we do not have to distinguish a single true causal variant among the variants constituting a single effect, but rather we need to consider them as a joint single haplotype effect (**Figure 3c**).

We asked whether such scenario is pervasive for *eSNP*s within the transcribed regions. First, we observed that the number of candidate causal variants with PIP>0.05 within the 95% credible set was significantly larger for *eSNP*s within the transcribed regions than those in other annotations such as promoters and enhancers (two-sided *t*-test *P*=4.6×10^-17^; **Figure 3d**). Second, we examined LD scores, which capture the weighted averaged number of SNPs in linkage with a lead SNP. We observed that *eSNP*s within the transcribed regions had significantly larger LD scores than promoters and enhancers (**Figure 3e**; **Methods**). We replicated this trend by using fine-mapped variants of independent bulk-eQTL data from GTEx^6^ with greater statistical significance (**Supplementary Figure 7**). These observations may support our proposed scenario that multiple causal variants in the transcribed regions within a haplotype tend to synergistically affect the target RNA’s structure, leading to differences in RNA abundance. In this scenario, all variants within a single effect can be causal, as opposed to variants within promoters or enhancers which span only ∼250 bp^55,56^ and most likely harbor one functional variant.

### Disease colocalization for nucleus- and cellular-specific

Finally, we investigated whether the eQTL localizing to certain subcellular compartments shared causal variants with disease GWAS, which helps us understand molecular mechanisms. Recently, there is growing discussion that GWAS loci are not sufficiently explained by putative causal variants from *cis*-eQTL studies. Mostafavi et al.^57^ proposed that the limited overlap might reflect systematic differences in causal variant locations and their target genes; disease GWAS variants are more likely to be under selective pressure and hence distribute further from TSS in the genes with high selective constraint than eQTL variants. Motivated by our observation of distinct causal variant locations between n- and c-eQTL, we tested each of them for colocalization with disease GWAS. Across 11 neuropsychiatric disorders (**Supplementary Table 1**; **Methods**), we identified 156 unique disease-*eGene* pairs that share the same causal variants (**Supplementary Table 2**). Among them, 42 and 77 disease-*eGene* pairs specifically colocalized with n-eQTL and c-eQTL, respectively, while 37 disease colocalizations were shared among both eQTL. For example, we identified n-eQTL in tubulin gamma complex component 4 (*TUBGCP4*) that colocalized with schizophrenia GWAS (lead SNP *P*=1.4×10^-14^, PP(H4_GWAS_)=0.90; **Figure 4a**). This locus had distinct causal variants at more powered c-eQTL (PP(H3_eQTL_)=0.94) with less significance (lead SNP *P*=4.2×10^-7^) that did not colocalize with schizophrenia (PP(H4_GWAS_)=5.0×10^-4^), suggesting that bulk eQTL had not sufficiently explained this locus. In another example, c-eQTL in the calcium voltage-gated channel subunit alpha1 I (*CACNA1I*) shared causal variants with schizophrenia GWAS with PP(H4_GWAS_)=0.97 (**Figure 4b**). In this locus, there were no shared causal variants (PP(H4_GWAS_)=0.10) between n-eQTL and schizophrenia. For such loci, the brain snRNA-seq or multiome RNA/ATAC-seq datasets^58–61^ might not adequately identify causal mechanisms. Taken together, we argue that subcellar localization of RNA matters in identifying molecular mechanisms of GWAS loci in addition to cell types, tissues, and conditions of eQTL.

**Figure 4.**
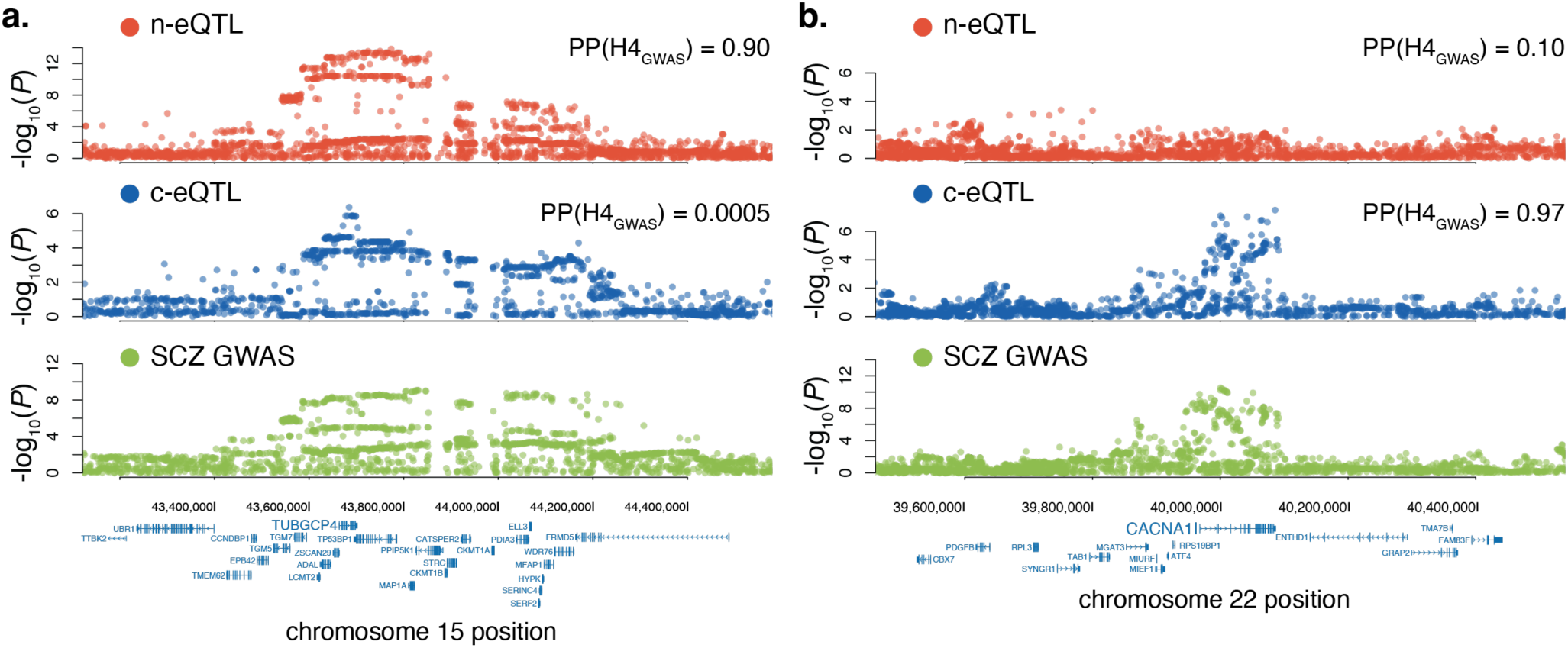
Subcellular-eQTL-specific disease colocalizations. Regional plots for eQTL and GWAS statistics (schizophrenia) at *TUBGCP4* (**a**) and *CACNA1* at locus (**b**). For both panels, top red plots show n-eQTL statistics, middle blue plots show c-eQTL statistics, and bottom green plots show GWAS summary statistics for schizophrenia. Posterior colocalization probability (H4: shared causal variant) was assessed between eQTL and GWAS.

## Discussion

In this study, we showed that the genetic variants affecting inter-individual differences in RNA abundance are, in many instances, entirely different between nuclear RNA and cellular RNA. In two different tissues, we argue that causal variants that are unique for n-eQTL are more likely to be driven by distal regulatory elements affecting transcriptional activity. In contrast, c-eQTL variants localized to transcribed regions and RBP binding sites, likely acting through altered post-transcriptional regulation.

Our data argues further argues that c-eQTL may be driven by multiple variants in tight linkage and located on the same haplotype in the transcribed regions. Generally, in causal variant fine-mapping, investigators have considered the possibility that multiple causal variants in a single locus might individually affect different molecular processes^50^, and methods assume that these variants are unlinked by LD, enabling multi-variant fine-mapping. However, our c-eQTL cases suggest that it may be inappropriate to narrow down the list of causal variants in a single effect to disentangle the LD structure^62^. All variants in a causal haplotype could, in concert, affect inter-individual differences in RNA structure, stability or degradation rate.

We finally demonstrated that nuclear- or cytosolic-specific eQTL causal variants sometimes colocalize with disease variants. There has been a growing debate on the reasons why there has been limited overlap between currently available eQTL variants and GWAS variants for complex diseases^9,10^. Major reasons discussed so far have included tissue- and cell-type specific eQTL variants and context-dependent effects, such as cellular activation^63–66^. Our observation highlights the importance of expanding our eQTL catalog in terms of (1) subcellular localization of RNA and (2) processes that affect RNA abundance after the transcriptional regulation.

We were intrigued by the instances when genetic disease risk variants sometimes colocalize with variants affecting nuclear RNA abundance but not cellular aggregated abundance. Recent genomic studies also suggest that genetic variants affecting epigenomic changes in the nucleus, such as chromatin accessibility, more likely colocalize with disease alleles even in the absence of corresponding changes in cellular RNA abundance^67,69^. One possible explanation is that for those genes, the rate of RNA transcription in the nucleus is more associated with translation efficiency^68^, rather than the accumulated sum of RNA in the cytosol in a steady state. Nuclear eQTL variants may capture cells’ ability to dynamically respond to a changing environment. Indeed, it has been recently demonstrated that the transcription rate might be coupled with translation efficiency^70^. Another study showed that for stress-response genes, newly synthesized RNA, as opposed to RNA pre-existing the stress condition, preferentially escapes translational repression as a mechanism to rapidly adapt to environment.^71^ These recent findings are in line with functional significance of n-eQTL alleles that affect transcription rate and their potential role in diseases.

Our study has several potential limitations. First, we compared nuclear and cellular RNA abundance to infer distinct genetic regulator for nuclear and cytoplasmic subfraction of RNA. Multiple previous studies have confirmed that cytoplasmic RNA molecules comprised the majority of cellular RNA across human tissues^19^. However, this could have reduced our power for discovery and more direct and powerful comparison might be achieved by extracting cytoplasmic RNA, if in the future such technologies are applied widely with matched nuclear RNA profiling. Second, all the datasets we analyzed used 3’-end RNA-sequencing method which does not allow us to perform transcript-level analyses. To partly address this, we tested colocalization of n-eQTL and c-eQTL with published sQTL from the DLPFC^80^. We observed that colocalizing probability with sQTL did not significantly vary between n-eQTL and c-eQTL (paired *t*-test *P*=0.14; **Methods**). This is biologically plausible, since the splicing machinery acts before RNA molecules are exported out of the nucleus. If in the future 5’-end or full-transcript technologies are applied for both nuclear and cytoplasmic fractions, these datasets might reveal more precise effect on alternative splicing. Third, to functionally prove regulatory effects of the identified eQTL variants, especially on RNA structure and RBP binding capacity, we need experimental perturbation approaches accompanied by molecular profiling of RNAs.

In summary, we demonstrated distinct regulatory mechanisms affecting RNA abundance that localize in the nucleus or in the cytosol which encompassed various post-transcriptional processes beyond conventional transcriptional regulation. These mechanisms can sometimes explain missing molecular causes of diseases, expanding our view on gene regulation and their association with pathogenic processes.

## Data Availability

The publicly available datasets from ROSMAP (https://www.synapse.org/#!Synapse:syn31512863) and AMP RA/SLE (https://arkportal.synapse.org/Explore/Projects/DetailsPage?Project=SLE_PhaseII) can be accessed via Synapse under Data Usage Agreement.

## Code Availability

The computational scripts related to this manuscript are available at https://github.com/immunogenomics/EarlyLate_RNA_eQTL.

## Methods

### Datasets

This study was performed in accordance with protocols approved by the Brigham and Women’s Hospital.

We used two independent sources of data for the main analyses in the paper. The first dataset is snRNA-seq and bulk RNA-seq from dorsolateral prefrontal cortex (DLPFC) of the postmortem brain tissue samples from the ROSMAP cohort^20^. We used this dataset as a discovery dataset since it had both RNA and genotype data from > 400 individuals. At the time of death, 34% of participants were cognitively nonimpaired, 26% were mildly impaired and 40% had dementia. 68% were female and 63% fulfilled a pathological diagnosis of Alzheimer’s disease by the National Institutes of Health (NIH) Reagan Criteria^20^. We downloaded the raw count matrix of snRNA-seq and whole-genome sequence (WGS) genotype from 424 individuals^21^ from Synapse under agreement on Controlled Data Usage. Since raw data from bulk RNA-seq was not available at the time of our analysis, we downloaded the available summary statistics of eQTL from the original study^21^ from Synapse, instead of running eQTL mapping on our own. We also obtained a list of significant *eGenes* as defined in the study through communication with the authors of the paper.

The second dataset is snRNA-seq and scRNA-seq from biopsied kidney tissues from the AMP RA/SLE consortium^22^. We used this dataset as a replication dataset. In brief, as part of the AMP RA/SLE Phase 2 consortium patients, individuals >16 years old undergoing a clinically indicated kidney biopsy to evaluate proteinuria were enrolled if they met sufficient criteria of systemic lupus erythematosus (SLE) diagnosis based on the revised American College of Rheumatology or the Systemic Lupus Erythematosus International Collaborating Clinics classification criteria, along with healthy control individuals. We processed the scRNA and snRNA expression matrices and genotype data, the details of which will be described in the flagship paper (*manuscript in preparation*).

### Genotype quality control

For the downloaded WGS data from the brain dataset, we selected variants that were (i) annotated as ‘PASS’, (ii) biallelic, (iii) variant call rate > 0.95, (iv) minor allele frequency (MAF) > 0.01 and (v) *P*_HWE_ > 1.0×10^-6^. For the SNP-chip genotype data from the kidney dataset, we applied the same genotype quality control and imputation steps as described in previous studies from AMP RA/SLE consortium^72,73^. In brief, we genotyped donors by using Illumina Multi-Ethnic Genotyping Array. We retained samples and variants with (i) sample call rate > 0.99, (ii) variant call rate > 0.99, (iii) MAF > 0.01, and (iv) *P*_HWE_ > 1.0×10^-6^. We performed haplotype phasing with SHAPEIT2 software and performed whole-genome imputation by using minimac3 software with a reference panel of 1000 Genomes Project phase 3^74^. After imputation, we selected variants with imputation *Rsq* > 0.7 and MAF > 0.05 as post-imputation QC.

### eQTL mapping

For snRNA-seq and scRNA-seq of both the brain and the kidney datasets, we pseudo-bulked and normalized the expression matrix using post-QC cells from all the cell types to maximize the statistical power in eQTL discovery and fine-mapping. We aggregated and averaged the log-normalized expression values from a given donor for a given gene. We retained genes with non-zero expression sample rate > 0.2 and performed inverse normal transformation of the aggregated expression across samples. We then performed PEER correction^75^ of the gene expression by including known covariates: Sex, age at death (the age of sample collection), study cohort, postmortem interval, and 5 first genotype principal component (PC) values for the brain dataset and sex, age, and 5 first genotype PC values for the kidney dataset. We confirmed that the PEER factors for correction are not colinear each other.

We performed standard *cis*-eQTL mapping for the genes after QC by linear models between corrected gene expression and genotype dosages. We used FastQTL software (version 2.0)^76^ with default settings. In addition to obtaining nominal association statistics from the linear models, we used FastQTL’s beta approximation to compute permutation *P* values from 10,000 permutations. Finally, to adjust for multiple hypothesis testing across genes and define significant *eGenes*, we obtained Storey’s *q* values using qvalue package (version 2.30.0) in R (version 4.2.3). We used *q* values < 0.10 as the level of FDR significant *eGenes* for these analyses for snRNA-seq and scRNA-seq.

### Fine-mapping and colocalization

We used coloc software (version 5.2.3) ^26,27^ to perform fine-mapping and colocalization analyses between n-eQTL and c-eQTL. The fine-mapped posterior probability for causal variants were calculated from the approximate Bayesian Factor (ABF) analysis implemented in finemap.abf() function with default settings. Importantly, these coloc analyses assume that there is a single causal variant in the *cis*-region for a given gene. This setting has a good feature for the goal of our analyses that the sum of posterior inclusion probability (PIP) across the variants in *cis*-region should always sum up to one when we analyze significant *eGenes*, which makes the analyses comparable across different genes. In addition, we discussed potential multiple causal variant scenarios when multiple variants had very similar PIPs in the Main text. For the differential enrichment test that is going to be described, we adjusted raw PIPs by dividing them by sum of PIPs across the *cis*-variants such that sum of PIPs adds up to one, since we have already defined significant *eGenes* by FDR and we only analyzed significant *eGenes*. For the colocalization between n-eQTL and c-eQTL, we used coloc.abf() function with default settings and defined posterior probability (PP) for hypothesis 3 (H3) for loci with distinct causal variants between n-eQTL and c-eQTL and PP for hypothesis 4 (H4) for loci with shared causal variants between n-eQTL and c-eQTL. We ensured that the sum of PP for H3 and PP for H4 added up to one, since we had already selected loci for the significant eGenes for both n-eQTL and c-eQTL by our definition. We excluded any loci (*eGene*s) with less than 200 genetic variants shared between n-eQTL and c-eQTL.

### Analyses with PsychENCODE dataset

We sought to compare our observation of effect size correlation and colocalization between n-eQTL and c-eQTL by comparing two similarly sized c-eQTLs (bulk RNA-seq eQTLs) in human brain prefrontal cortex. The first dataset was bulk RNA-seq eQTL from the ROSMAP cohort (*n*=1,092) that we had described in the previous section, and the second dataset was bulk RNA-seq eQTL from the PsychENCODE Consortium. The PsychENCODE^28^ collected bulk RNA-seq and genotype data from 1,387 individuals and released eQTL summary statistics.

We assessed the lead SNP effect size correlation between ROSMAP and PsychENCODE. We took 4289 shared *eGene*s from both cohorts, aligned the effect alleles, intersected *cis*-genetic variants for each eGene shared between the two cohorts, and calculated the Pearson’s correlation *R* of the beta coefficients from the two cohorts by selecting one most significant genetic variant from the ROSMAP study for each of the 4289 shared *eGene*s. We next assessed colocalization of causal variants between ROSMAP c-eQTL and PsychENCODE c-eQTL. We ran coloc.abf() function for the shared *eGene*s between ROSMAP and PsychENCODE and only retained loci with at least 200 genetic variants shared between ROSMAP and PsychENCODE. Similarly to the primary analysis, we ensured that the sum of PP for H3 and PP for H4 added up to one, since we had already selected loci to be analyzed for the significant eGenes for both studies.

### Differential enrichment test

To quantify differential enrichment of causal eQTL variants between n-eQTL or c-eQTL within a specific genomic annotation or unit, we defined an enrichment statistic *t*_*pip*_ for each of the genomic units. For the genomic annotations or units to be tested for enrichment, we examined promoters, candidate cis-regulatory elements (cCREs) with varying distances from TSS, exonic and transcribed regions, 3’ and 5’ UTRs, RBP binding sites, and m^6^A modification sites. We defined exonic regions by aggregating all possible exonic regions across all patterns of transcripts for the target gene registered at RefGene database (GRCh37). The brain specific cCREs were based on Brain Open Chromatin Atlas^77^ and PsychENCODE enhancer list for prefrontal cortex ^28^. The kidney specific cCREs were based on cCRE annotation in the human kidney sample from ENCODE v3 (accession: ENCSR751YZB). We defined potential RBP binding sites by using peaks from by eCLIP assay released from ENCODE consortium^29^ for 168 unique RBPs for HepG2 and K562 cell lines as well as adrenal gland tissue. We defined m^6^A modification sites from published m^6^A -seq data from 60 Yoruba lymphoblast cell lines (LCLs).^17^ For RBP binding sites and m^6^A modification sites, we ensured that those regions are included in the transcribed region of the *eGene* which we tested for enrichment so that we have accurate insights into the post-transcriptional regulation for the target *eGene* but not for the other genes.

Based on fine-mapped PIPs for either n-eQTL or c-eQTL, we first took sum of PIPs for all the SNPs that belong to a given genomic unit *c* (e.g., promoter) for each eGene *g* as an enrichment statistics for {*nucleus or cell*} eQTLs:

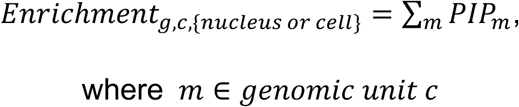

*m* indicates SNPs in cis-region for eGene *g* that belong to the genomic unit *c*.

We then compared *Enrichment_g_*_,*c*,*cell*_ and *Enrichment_g_*_,*c*,*nucleus*_ for each eGene *g*, and ran paired T-test across all the shared eGenes between n-eQTL or c-eQTL to calculate *t*_*pip*,*c*_ for the genomic unit *c*.

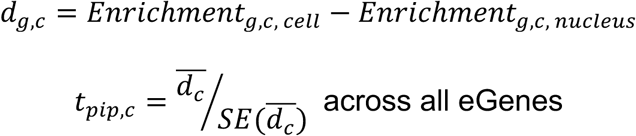

Given the subtraction direction above, the positive *t*_*pip*,*c*_ value indicates that the genomic unit *c* has stronger enrichment for c-eQTL causal variants, whereas the negative *t*_*pip*_, *c* value indicates that it has stronger enrichment for n-eQTL causal variants.

### Test for asymmetric distribution of causal eQTL variants

We selected variants with PIP > 0.7 as highly probable causal variant for each *eGene*. We obtained a list of such variants and their distance to transcription start site (TSS) with accounting for the gene’s strand. We concatenated the list across all the eGenes separately for n-eQTL and c-eQTL. We then binned the *cis*-region (+-1 Mb window from TSS) by a window size of 20 kb. Separately for n-eQTL and c-eQTL, we calculated the fraction of causal variants belonging to each bin as the proportion of SNPs belonging to the bin similarly to ref^57^. We also calculated the 95% confidence intervals (CIs) of these proportions by boostrapping the variants in the list by 1,000 times. To statistically test the differences in the proportion between n-eQTL and c-eQTL for each bin, we performed chi-squared test for two-by-two table, with the number of causal variants within the bin and outside of the bin (columns) for n-eQTL and c-eQTL (rows).

We also tested asymmetric distribution of causal variants relative to the TSS as the center of the distribution. We defined the skewness statistics^33,34^ as follows using 3^rd^ moment of the signed distances between the causal variant and the TSS.

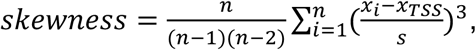

where *n* denotes the total number of causal variants. *x*_*i*_ indicates the genomic position for SNP *i*, *x*_*TSS*_ indicates the genomic position for TSS of the *eGene*, and *s* indicates standard deviation of the signed distances (i.e., differences in genomic positions) between the variant and the TSS. We flipped the sign of *x*_*i*_ − *x*_*TSS*_ if the *eGene* is coded in the negative strand.

### Baseline enrichment test

We also assessed the baseline distribution of the causal variants for the genomic units in n-eQTL and c-eQTL (**Supplementary Figure 3**). To compare enrichment across the genomic units rather than within a unit as we did in the *t*_*pip*_ analyses, we needed to adjust for the length and the number of variants within the genomic units and across the *cis*-region.

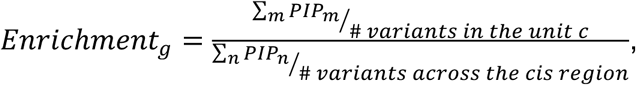

where *m* ∈ *causal variants in genomic unit c*,

and *n* ∈ *causal variants in the entire cis region*. We defined the causal variants with variants with PIPs > {0.3, 0.5, 0.7}. We then calculated the overall baseline enrichment value by taking the mean of *Enrichment_g_* across all *n* eGenes,

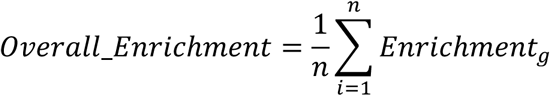

and derived the 95% CIs by bootstrappoing genes by 1,000 times.

### Conditional enrichment test

In assessing *t*_*pip*_ in genomic annotations included in the transcribed regions in principle (i.e., UTRs, exons, introns, and m^6^A modification sites and RBP binding sites), we wanted to account for known strong enrichment within the transcribed regions. To this end, we performed conditional analyses for these genomic units (**Supplementary Figure 4**). We first defined *Fractional*_*Enrichment* statistics by dividing sum of PIPs in a given genomic unit *c* by sum of PIPs in the transcribed regions as follows:

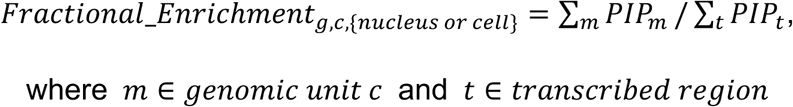

We also ensured that all SNPs *m* that belong to the genomic unit *c* are also within the transcribed region of of the gene *g*. Similarly to the primary analyses, we then compared *Fractional*_*Enrichment_g_*_,*c*,*cell*_ and *Fractional*_*Enrichment_g_*_,*c*,*nucleus*_ for each eGene *g*, and ran paired T-test across all the shared eGenes between n-eQTL or c-eQTL to calculate *conditional*_*t*_*pip*,*c*_ for the genomic unit *c*.

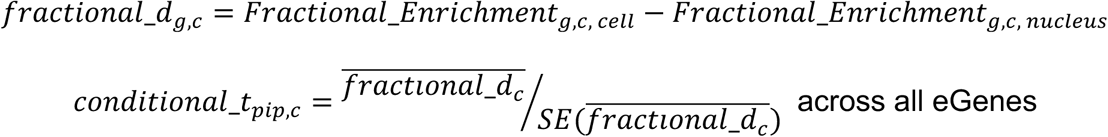

### Credible set size

We assessed significant differences in the size fine-mapped credible set between *eGene*s with probable causal variants within transcribed regions or *eGene*s with probable causal variants within conventional promoters or cCREs. For each eGene *g* in c-eQTL in the brain dataset, we assessed the genomic unit with the highest *Enrichment_g_*_,*c*, *cell*_ with assuring the *Enrichment_g_*_,*c*, *cell*_ > 0.05. Based on this, we categorized *eGene*s into promoter-driven, enhancer (cCREs)-driven, transcribed region-driven, and other ones. We constructed 95% credible sets for eGenes by sorting variants based on their PIPs in a descending order and cumulatively summing and collecting them until the cumulative sum is 0.95. We note that we used variants with at least PIP > 0.05 for this analysis (i.e., maximum credible set size is 19) since we intended to avoid loci (*eGene*s) with hundreds of uncertain variants with very low PIPs. We then compared the number of variants within the 95% credible sets between transcribed region driven eGenes and promoter- or enhancer-driven *eGene*s by T test.

### Tagged variants in LD

We also assessed significant differences in the number of genetic variants tagged by LD between probable causal variants in transcribed region-driven *eGene*s and those in promoter- or enhancer-driven *eGene*s. Using the WGS genotype data in the brain dataset, we comprehensively calculated the LD *R*^2^ between a sentinel variant with maximum PIP (> 0.05) for a given *eGene* and all other variants in *cis*-region for the *eGene* within 1 Mb window. We defined the LD scores for each *eGene* by summing the total LD *R*^2^ of variants that had greater values of LD *R*^2^ than a sliding LD threshold {0,0.1,0.3,0.5,0.7} to assess the effect of removing noisy *R*^2^ from weakly tagged variants. We associated these LD scores from each LD threshold with the categorical variable whether the *eGene* is classified as transcribed region-driven or promoter- or enhancer-driven, with including MAF of the sentinel variant as a covariate.

### Analyses with GTEx dataset

To examine if the observation on credible set size and tagged variants in our brain dataset is generalizable across other tissues’ eQTL, we used published GTEx v8 dataset. We used GTEx fine-mapping results by CAVIAR. Similarly to our primary analysis in the brain dataset, we defined promoter-driven, enhancer (cCREs)-driven, and transcribed region-driven *eGene*s for each tissue. cCRE regions were defined using cCRE annotation from ENCODE consortium^78^ that covers many human cell lines and tissues. We ran the same test for comparison of the 95% credible set size between transcribed region-driven eGenes and promoter- or enhancer-driven eGenes by T test. We also ran the same association test between the LD scores for the sentinel variant of an *eGene* and the categorical variable whether the *eGene* is classified as transcribed region-driven or promoter- or enhancer-driven, with including MAF of the sentinel variant as a covariate. We used 1000 Genomes Project phase 3 genotype data to calculate the LD scores and MAF.

### Disease colocalization

Using the brain dataset, we conducted colocalization analysis of causal variants from n-eQTL or c-eQTL and those from neuropsychiatric diseases. We downloaded publicly available GWAS summary statistics of 10 neuropsychiatric disorders from the Psychiatric Genomics Consortium (Alzheimer’s disease, anorexia nervosa, attention deficit hyperactivity disorder, autism spectrum disorder, bipolar disorder, major depressive disorder, obsessive compulsive symptoms, panic disorder, post-traumatic stress disorder, schizophrenia) and one from GWAS catalog (Parkinson’s disease^79^; **Supplementary Table 1** for details). We defined loci for colocalization based on the *cis*-regions for all eGenes from either n-eQTL or c-eQTL in the brain dataset. For each disease GWAS, if the *cis*-region contains any variant with genome-wide significance (*P*_GWAS_ < 5×10^-8^), we proceeded on assessing colocalization of causal variants between eQTL and GWAS by using coloc software with default settings. If they do not contain any genome-wide significant variant, we did not perform colocalization analyses. We also excluded any loci with less than 200 genetic variants shared between eQTL and GWAS.

### sQTL colocalization

We obtained sQTL summary statistics from 450 DLPFC samples in ROSMAP through personal communication with the author on the original publication^80^. We selected 1,313 genes and their trascripts that are significant in previously described n-eQTL and c-eQTL, as well as in the sQTL study. We tested colocalization of causal variants between sQTL and n-eQTL and between sQTL and c-eQTL by using coloc with the same procedures described in the previous sections. We finally tested whether there is a statistically significant difference in the posterior probability of shared causal variants (PP(H4) for sQTL and n-eQTL vs. PP(H4) for sQTL and c-eQTL) by paired *t*-test across 1,313 genes.

## Acknowledgments

We would like to thank Phoenix Mu, Benjamin Strober, Alkes Price, Shamil Sunyaev, and Sasha Gusev for helpful discussions. We thank Joseph Mears, Thomas Eisenhaure, Qian Xiao, Sid Gurajala, Betty Diamond, Nir Hacohen, Anne Davidson, and Deepak Rao for construction and analyses of AMP RA/SLE Network dataset. We thank Towfique Raj for providing the splicing QTL summary statistics. This work is supported in part by funding from the National Institutes of Health (R01AR063759, U01HG012009, UC2AR081023). The results published here are in whole or in part based on data obtained from the ARK Portal (https://arkportal.synapse.org/). This work was supported by the Accelerating Medicines Partnership® Rheumatoid Arthritis and Systemic Lupus Erythematosus (AMP® RA/SLE) Program. AMP® is a public-private partnership (AbbVie Inc., Arthritis Foundation, Bristol-Myers Squibb Company, Foundation for the National Institutes of Health, GlaxoSmithKline, Janssen Research and Development, LLC, Lupus Foundation of America, Lupus Research Alliance, Merck & Co., Inc., National Institute of Allergy and Infectious Diseases, National Institute of Arthritis and Musculoskeletal and Skin Diseases, Pfizer Inc., Rheumatology Research Foundation, Sanofi and Takeda Pharmaceuticals International, Inc.) created to develop new ways of identifying and validating promising biological targets for diagnostics and drug development Funding was provided through grants from the National Institutes of Health (UH2-AR067676, UH2-AR067677, UH2-AR067679, UH2-AR067681, UH2-AR067685, UH2-AR067688, UH2-AR067689, UH2-AR067690, UH2-AR067691, UH2-AR067694, and UM2-AR067678).

## Author Contributions

S.S. and S.R. conceived the work and wrote the manuscript. S.S. conducted all analyses with inputs from S.R. Accelerating Medicines Partnership®: RA/SLE Network provided the kidney dataset.

