## Supplementary Figures for "Early and late RNA eQTL are driven by different genetic mechanisms"

Sakaue et al.

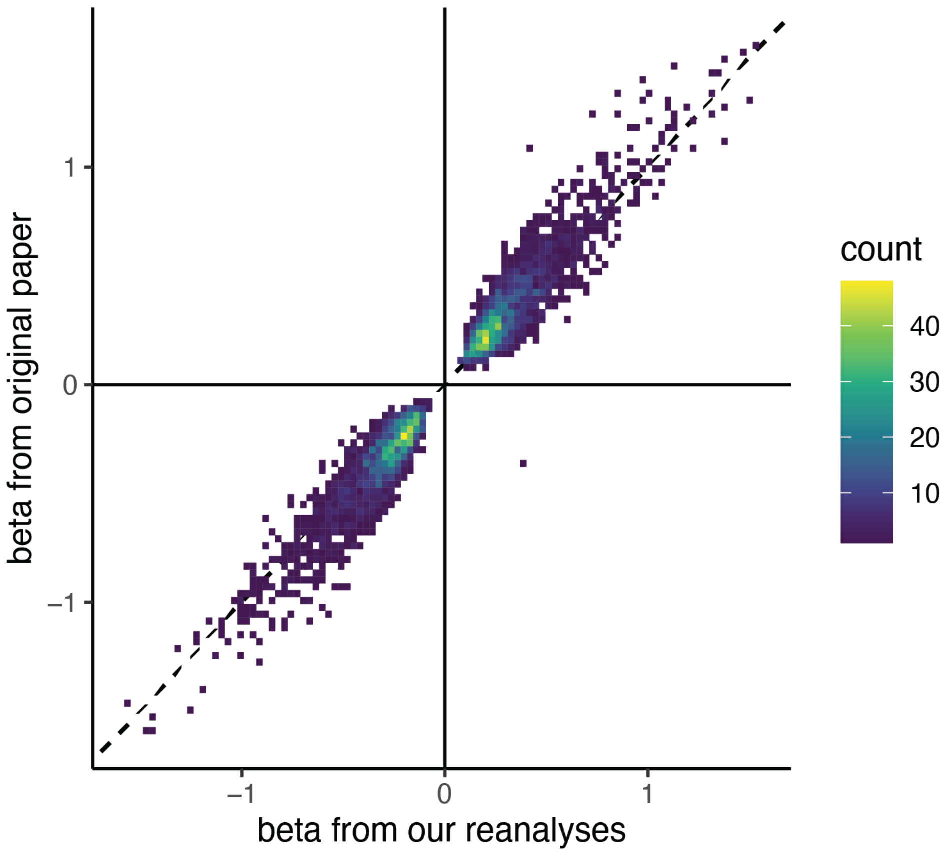


**Supplementary Figure 1 I Correlation of eQTL effect estimates between our analyses and the original publication.**

We show the consistency of eQTL effect estimates in excitatory neurons between our re-analyses from downloaded datasets (x-axis) and summary statistics from their original publication (y-axis). Each point originally was each significant eSNP-eGene pairs in both our study and their study, and eSNP was chosen from the lead SNPs from our study. For visibility, then these individual points were aggregated into bins to show the density of observations as shown in the right color scale, using geom_bin2d package in ggplot2 in R.

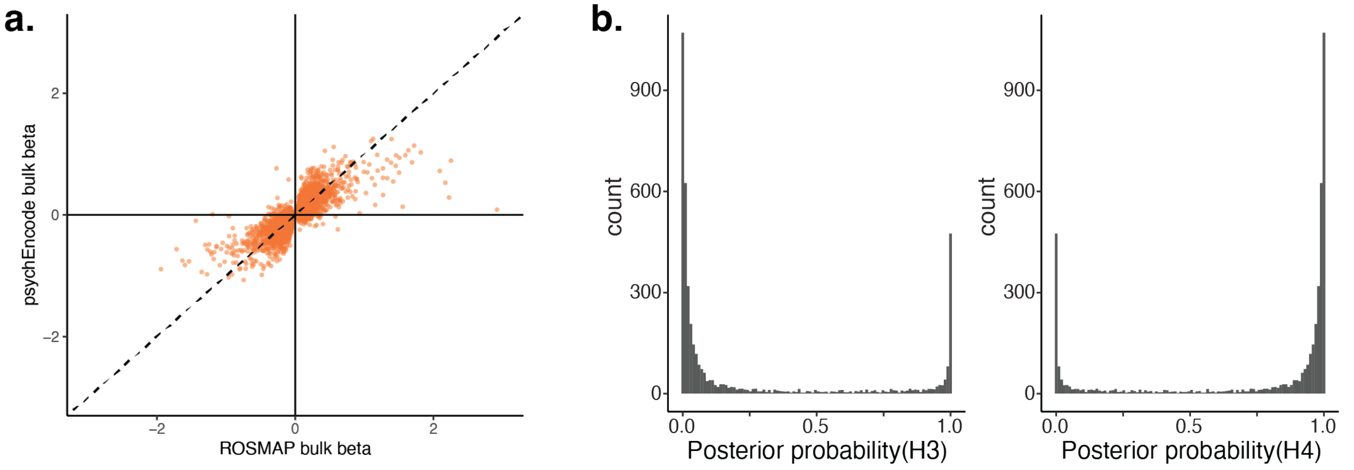


**Supplementary Figure 2 | Comparison of bulk RNA-seq eQTL statistics between ROSMAP and psychENCODE consortia.**

**a**. Scatter plot for beta coefficients for the same eSNP-eGene pairs between ROSMAP bulk RNA-seq eQTL (x-axis) and psychENCODE bulk RNA-seq eQTL (y-axis) in the brain dataset. Each dot represents one eSNP-eGene pair, and eSNPs were selected based on n-eQTL statistics. **b**. Histogram of coloc posterior colocalization probability for H3 (left; distinct causal variant between ROSMAP bulk RNA-seq eQTL and psychENCODE bulk RNA-seq eQTL) and H4 (right; shared causal variant between ROSMAP bulk RNA-seq eQTL and psychENCODE bulk RNA-seq eQTL).

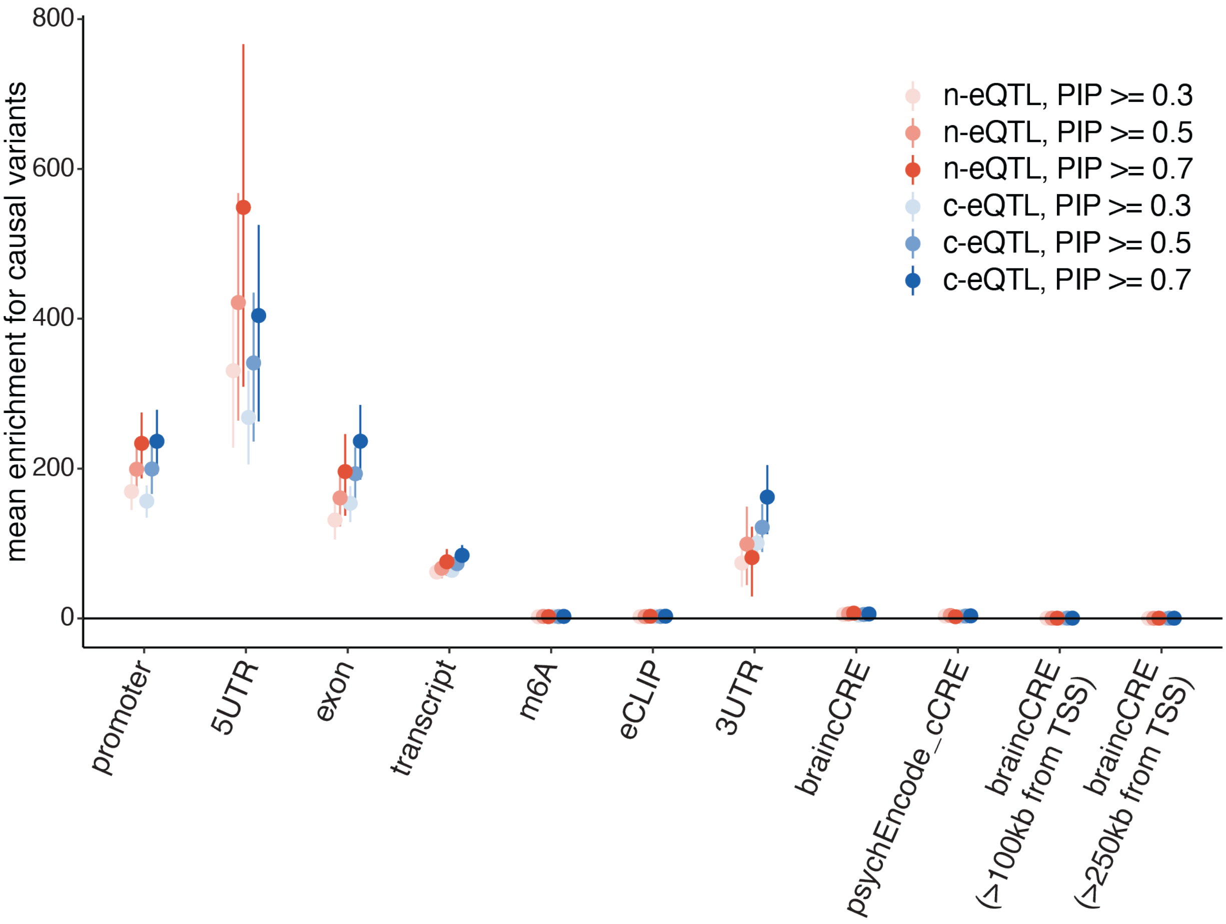


**Supplementary Figure 3 I Enrichment of candidate causal variants within genomic features.**

We assessed baseline enrichment of causal variants in a given genomic unit that we tested for differential enrichment between n-eQTL and c-eQTL. We defined the baseline enrichment (the means across genes as dots and 95% confidence intervals by bootstrapping genes as bars) as: (sum of PIPs for causal variants with PIP >= threshold within a genomic unit / # variants tested within a genomic unit) / (sum of PIPs for causal variants with PIP >= threshold within cis-region / # variants tested within cis region). The threshold to define PIPs were chosen from {0.3, 0.5, 0.7} and colored differently as shown in the color legend in the figure. We note that in each of the PIP threshold, we excluded any loci with no causal variants above the PIP threshold.

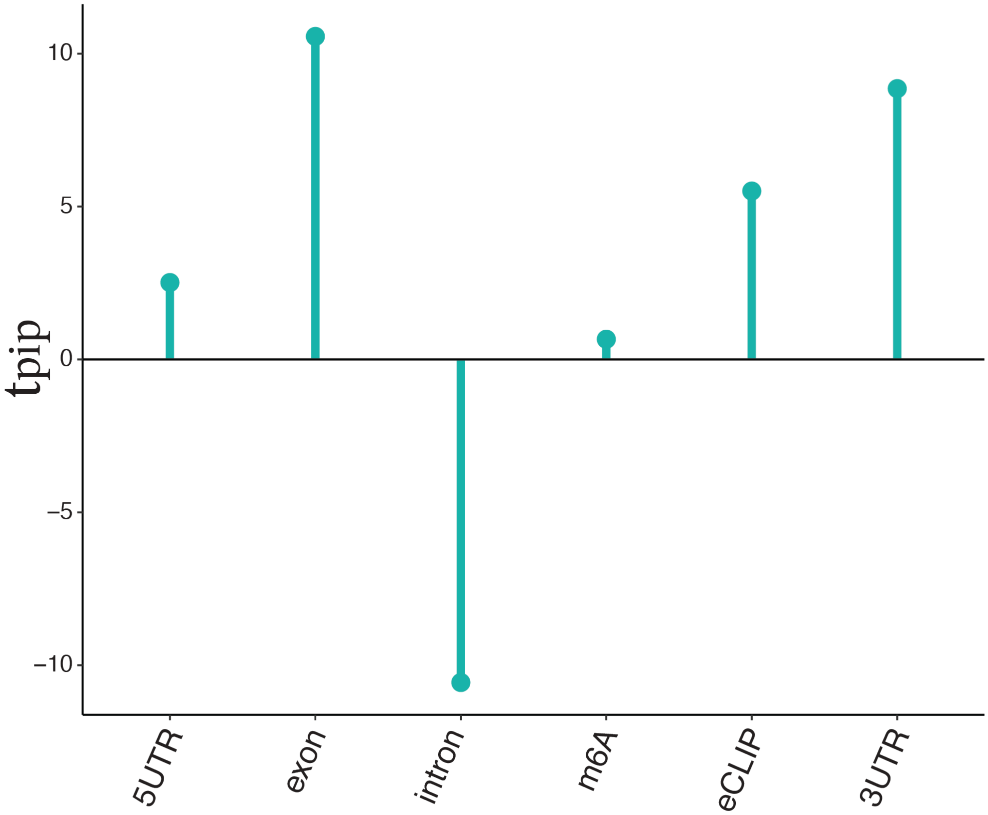


**Supplementary Figure 4 I Conditional analyses adjusting for transcribed regions for differential enrichment for causal variants between c-eQTL and n-eQTL in the brain dataset.**
$t_{pip}$ statistics in the brain data after adjusting for enrichment of causal variants within the transcribed regions. We only selected genomic regions for each category (x-axis) within the transcribed regions, and derived conditional enrichment by dividing sum of PIPs in the category by sum of PIPs in the transcribed regions (see **Methods**).

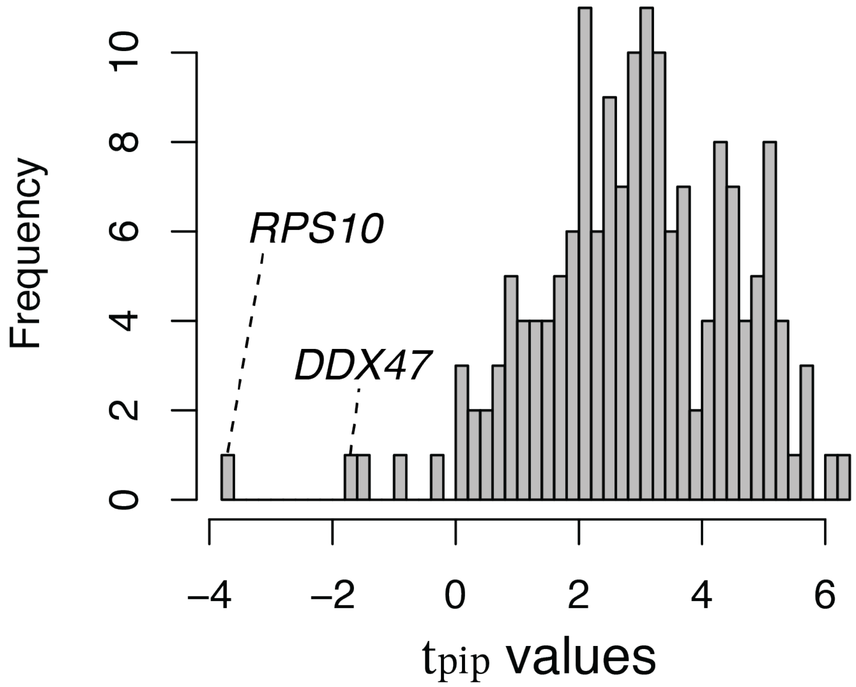


**Supplementary Figure 5 I RNA binding protein specific differential enrichment for causal variants between c-eQTL and n-eQTL.**

A histogram showing the $t_{pip}$ enrichment statistics comparing enrichment of c-eQTL and n-eQTL causal variant locations within each of the RNA binding protein (RBP)-specific eCLIP datasets in the brain dataset (see **Main Text** and **Methods**). Two genes are annotated for nominally significant negative $t_{pip}$ RBPs.

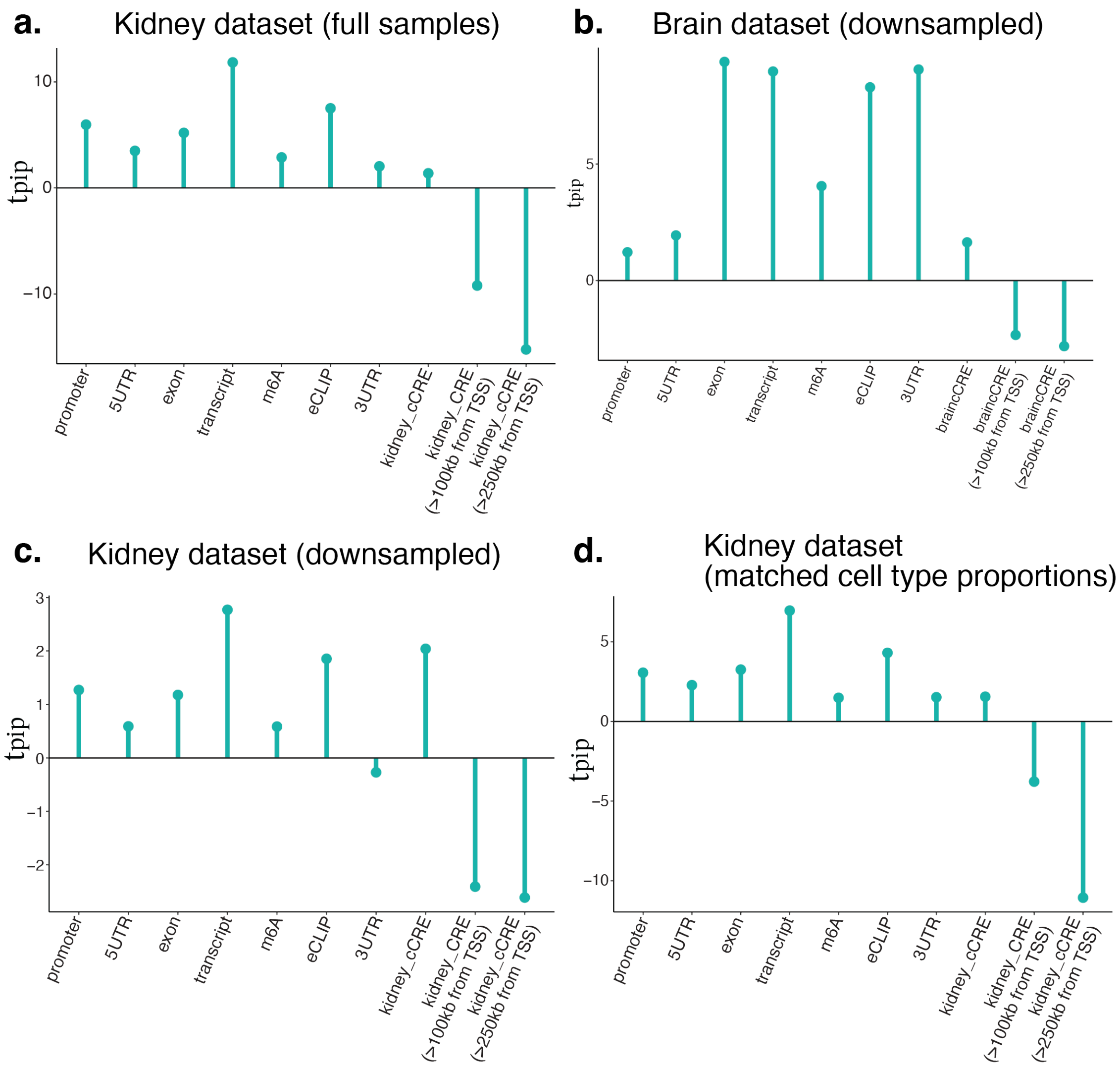


**Supplementary Figure 6 I Replication and sensitivity analyses for differential enrichment for causal variants between c-eQTL and n-eQTL.**

$t_{pip}$ statistics in four different datasets comparing enrichment of c-eQTL and n-eQTL causal variant locations are shown to replicate the finding observed in the primary brain dataset (**Figure 2c**). **a**. Kidney dataset with full samples of snRNA-seq and scRNA-seq. b. Brain dataset with simulated downsampling in bulk RNA-seq at the summary statistics level to have the same number of individuals between snRNA-seq and bulk RNA-seq. **c**. Kidney dataset with downsampling of scRNA-seq so that the snRNA-seq and scRNA-seq data are derived from the exactly matched individuals. **d**. Kidney dataset with randomly sampled cells to have matched proportion of cell types between snRNA-seq and scRNA-seq.

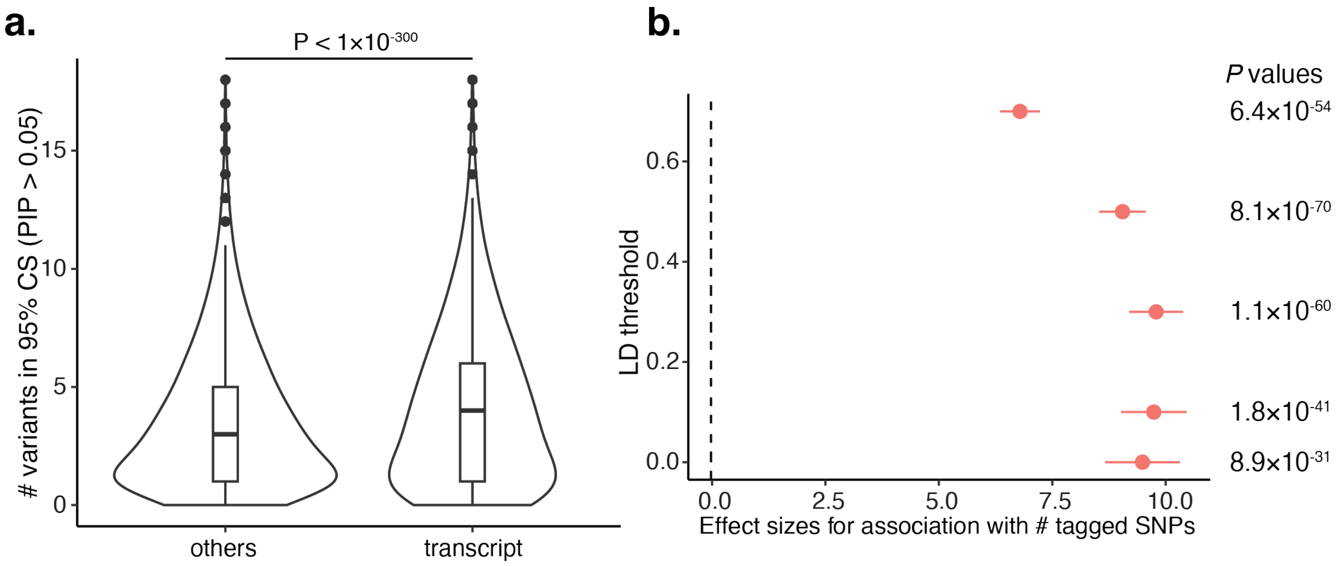


**Supplementary Figure 7 I The number of tagged variants in linkage disequilibrium and the number of variants in credible sets in GTEx fine-mapping resuls.**
This is a replication analysis of **Figure 3d** and **3e** in the **Main Text**, using an independent GTEx dataset. **a**. Violin plots indicating the distribution of the number of variants with PIP > 0.05 within 95% credible sets for eGenes with causal variants within transcribed regions (right) and all the other eGenes (left). **b**. Beta coefficients (circles; x-axis) and standard error (bars) from the association between the number of tag SNPs in each LD $R^2$ threshold (y-axis) and whether the eSNP is in the transcribed region or not (see **Main Text** and **Methods** for details).
